# mBARq: a versatile and user-friendly framework for the analysis of DNA barcodes from transposon insertion libraries, knockout mutants and isogenic strain populations

**DOI:** 10.1101/2023.11.27.568830

**Authors:** Anna Sintsova, Hans-Joachim Ruscheweyh, Christopher M. Field, Lilith Feer, Bidong D. Nguyen, Benjamin Daniel, Wolf-Dietrich Hardt, Julia A. Vorholt, Shinichi Sunagawa

## Abstract

DNA barcoding has become a powerful tool for assessing the fitness of strains in a variety of studies, including random transposon mutagenesis screens, attenuation of site-directed mutants, and population dynamics of isogenic strain pools. However, the statistical analysis, visualization and contextualization of the data resulting from such experiments can be complex and require bioinformatic skills. Here, we developed mBARq, a user-friendly tool designed to simplify these steps for diverse experimental setups. The tool is seamlessly integrated with an intuitive web app for interactive data exploration via the STRING and KEGG databases to accelerate scientific discovery.

## BACKGROUND

Advances in DNA sequencing and computational technologies have facilitated the assembly of millions of microbial genomes, leading to the continuous discovery of new genes [1]. However, the characterization of these genes has not kept pace with the rate of their discovery, since gathering functional insights, for example through individual gene knockouts, remains slow and labor-intensive. As a result, many genes remain uncharacterized, even in extensively studied organisms [2]. To fully explore the microbially encoded sequence space, it is crucial to increase the throughput of testing the effect of individual genes on cell growth and reproduction (i.e., fitness). However, traditional methods, such as comparing the fitness between individual mutant and wild type strains under specific conditions, do not scale well when dealing with a large number of genes and conditions.

A more effective alternative to such traditional approaches is to analyze multiple mutants in the same experiment. One powerful method to bridge the sequence-to-function gap is transposon-based mutagenesis coupled to next-generation sequencing, or transposon-insertion sequencing (TIS). TIS identifies genomic loci that contribute to organismal fitness under different experimental conditions [3]. Recent advances in the conventional TIS protocol, such as including a random DNA barcode sequence into each transposon for screening by PCR [3–6], have significantly increased experimental throughput. This approach (i.e., random barcode transposon mutagenesis coupled with sequencing, RB-TnSeq) is increasingly employed to study fitness effects of genes, as well as to improve the annotation of uncharacterized protein families across diverse bacterial species [6]. This methodology has also been applied to elucidate genotype-phenotype relationships in eukaryotic model systems [7–9]. However, while the experimental methodology has been well established, the processing and analysis of the resulting data remain a challenge.

RB-TnSeq data analysis involves several steps, including identifying the chromosomal positions of barcoded insertions (mapping), quantifying the barcoded strains across experimental conditions (counting), and ultimately, identifying the fitness factors or genomic loci whose functions are essential for or affect the reproduction of organisms in a given experimental setting (statistical analysis). However, each of these steps has to be adapted to a specific study, as there is great variation in experimental protocols, in library creation and in experimental design [4–6,9]. While a rich ecosystem of analysis and visualization tools exists for TIS [3], the only code available for the analysis of RB-TnSeq data is a collection of scripts that accompanied the original publication [5], and it is unclear whether the code can be adapted to RB-TnSeq libraries generated by other experimental protocols. Furthermore, a consensus on statistical procedures for the analysis of RB-TnSeq data is still lacking, and user-friendly, interactive tools are needed to contextualize the results and reduce the dependency on bioinformatic expertise.

In addition to studying the fitness effect or the essentiality of genes, barcoded sequencing data have also been used to study the dynamics of strain populations. In this approach, known barcodes are introduced at neutral genomic loci to trace isogenic strains over time and/or space. These barcoded strains are used to understand microbial population evolutionary trajectories, colonization bottlenecks, rates of immigration into a new niche, death and replication rates, and priority effects [10]. However, current tools generally lack the flexibility to analyze barcoded sequencing data from these types of experiments, or have optimized only specific parts of the analysis, such as barcode clustering accuracy [11], and lack the required mapping and analysis capabilities required for RB-TnSeq experiments.

Here, we introduce mBARq (pronounced: ‘embark’), a versatile and user-friendly framework for the analysis and interpretation of RB-TnSeq and other barcoded sequencing data. A command line tool allows mapping, counting and statistical analysis of RB-TnSeq data. Notably, we adapted a novel statistical framework [12,13] and benchmarked it using experimentally validated data [4] to show that it results in higher sensitivity, while retaining similar precision to previously published methods. In addition, we demonstrate that mBARq can also be applied to the analysis of barcoded isogenic strains to investigate their population dynamics. Finally, a companion web app enables customized quality control, visualization of the results and exploratory data analysis via integration with the STRING [14] and KEGG [15] databases.

## RESULTS

### Implementation of a versatile tool and web app for the analysis of barcoded sequencing data

We aimed to develop a versatile tool for the profiling and analysis of barcoded transposon-insertion libraries. Here, we describe the workflow applicable to RB-TnSeq experiments, with modifications for other experimental setups discussed later. At each step, seamless integration with a user-friendly web app allows non-experts to intuitively explore the results of individual steps.

#### Mapping

The first steps of a RB-TnSeq experiment consist of creating a transposon mutant library (for details see [3–5]), identifying the insertion site of each transposon, and matching the unique barcode that is linked to each transposon to its insertion site (Fig. 1A, panels i and ii). Experimentally, once the library is created, the barcoded transposons and adjacent host DNA are PCR-amplified, by combining a transposon-specific and random primer, and sequenced (Fig. 1A, panel i). Following the sequencing, mBARq uses quality-controlled sequencing data together with a genome sequence (FASTA format) and annotation file (GFF format) for the organism of interest to generate a library map, *i.e.,* a table, reporting the specific genomic position of each insertion, as well as any overlapping features of interest (gene, CDS, *etc.*) (Fig. 1A, panel ii). The genome sequence and annotation files can be either downloaded from public databases, for already sequenced genomes, or generated by the researcher, when investigating novel strains. The researcher can upload the library map to the mBARq web app to interactively assess the insertion coverage of transposons across the genome and generate summary statistics for the library (Fig. 1A, panel iii).

**Figure 1:**
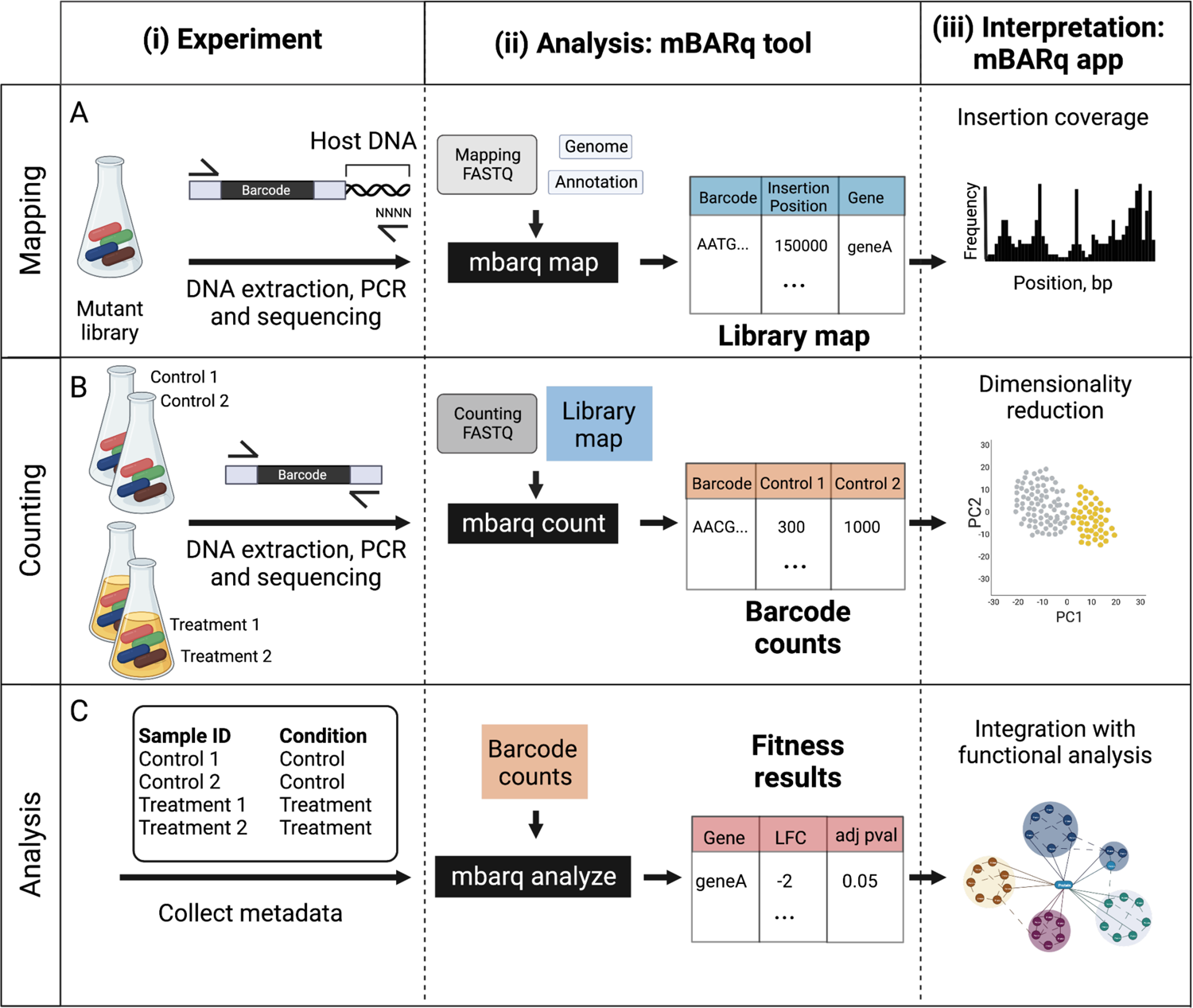
A universal and versatile framework for the analysis of barcoded transposon mutagenesis screens. **A**. Mapping. The mapping step determines the genomic location of each of the barcoded insertions in the mutant library. On the experimental side, this involves (i) extracting DNA from the mutant library, amplifying and sequencing the barcode, as well as a stretch of the host chromosome. The amplification in this step is accomplished using a PCR reaction with a construct-specific primer, and a random primer to allow host amplification. (ii) Using the sequencing data generated in (i), the mBARq tool generates a library map, which specifies the genomic position for each barcoded insertion, as well as genomic features associated with it. (iii) Users can upload the library map generated in (ii) to the mBARq web app to visualize insertion coverage across the genome, and generate library statistics (*i.e*., number of unique insertions, number of genes with an insertion, *etc.*). **B**. Counting. The experimental setup for the transposon mutagenesis screen involves (i) subjecting mutant libraries to a specific challenge (i.e., drug treatment, specific culture conditions). This challenge is followed by DNA extraction, barcode amplification and sequencing steps for each of the samples. The amplification in this case is accomplished using two construct specific primers. (ii) Using the sequencing data from this step and the library map created in A, the mBARq tool quantifies the abundance of each of the barcodes across samples and generates a barcode count table. (iii) Users can upload the barcode count table generated in (ii) to the mBARq web app for interactive exploratory data analysis. **C**. Statistical analysis. (i) mBARq allows the user to identify which mutants were sensitive to the challenge administered in B. This is accomplished by quantifying the differences in abundances of barcodes associated with each gene before and after the challenge. Using the metadata about the experiment (i) and barcode counts generated in B, the mBARq tool can perform statistical analysis of the barcode abundances to provide a fitness results table, listing log_2_ fold changes (LFC) and statistics for each gene that was disrupted in the library (ii). (iii) Users can upload the fitness results table generated in (ii) to the mBARq web app for functional analysis with STRING and KEGG databases. Figure created with BioRender.com.

#### Counting

In a typical RB-TnSeq experiment, the barcoded mutant library is subjected to a challenge (e.g., drug treatment, culture on a specific carbon source, or an *in vivo* pathogenesis model) to identify condition-specific fitness factors (Fig. 1B, panel i). After the challenge, a sequencing library is generated for each sample, i.e., the input pools (libraries before the challenge) and output pools (libraries after the challenge), by PCR using primers targeting the random barcodes. To quantify the abundance of each mutant in the input and the output pools, mBARq uses the raw sequencing data (FASTQ format) generated from this step, alongside the library map (Fig. 1B, panel ii). This results in a barcode count table listing the abundance of each barcode for the input sample and for each of the output samples. This count table can be uploaded to the mBARq web app to create an annotated principal component analysis (PCA) plot and to explore the barcode abundances for any gene of interest (Fig. 1B, panel iii).

By default, mBARq is set up to process Tn5-generated libraries with 17 bp barcodes. However, the entire workflow can be customized to the specific transposon used for library generation to ensure broad applicability of mBARq to diverse library construction methods. More detailed documentation and explanation of the steps described above are available online [16].

#### Statistical analysis

Statistical analysis of the count table allows the identification of condition-specific fitness factors, *i.e.*, genes whose loss negatively affects organismal growth. The analysis step implemented in mBARq allows for comparisons between two experimental conditions (control/treatment) using a robust statistical framework (MAGeCK [12,13]). While it was originally developed for pooled CRISPR knockout screens, we adapted this method for mBARq due to its demonstrated algorithmic advantages [17] and the methodological similarities between RB-TnSeq and CRISPR knockout screen data. Specifically, the MAGeCK framework addresses difficulties in estimating read counts with a small number of replicates and varying effects of different barcode insertion sites on gene fitness [12]. It also aggregates information from multiple insertions into the same gene. Extending MAGeCK’s functionality, mBARq allows for the incorporation of wild type isogenic controls for data quality control and normalization, which is recommended for *in vivo* experiments to account for potential population bottlenecks [4,10,18].

The mBARq analysis step requires a previously generated barcode count table and metadata specifying the condition for each of the samples (e.g., treatment or control; Fig. 1C, panel i). Running the analysis step produces the results table that includes log_2_ fold changes (LFC) and adjusted p-values for each gene in the library (Fig. 1C, panel ii). This results table can then be transferred to the mBARq web app (Fig. 1C, panel iii) to prioritize hits. User-defined genes of interest, based on LFC and adjusted p-value thresholds, can be subsequently analyzed using a STRING database [14]. Uploading the results to the STRING database allows for functional enrichment analyses and may reveal protein-protein interactions (PPIs) among the identified fitness factors or with other proteins. Additionally, for organisms present in the Kyoto Encyclopedia of Genes and Genomes (KEGG) database [15], it is possible to superimpose the results onto KEGG metabolic maps to obtain an integrated view of the results and facilitate data interpretation.

In the following sections, we demonstrate the flexibility, sensitivity, precision, and visualization capabilities of the mBARq framework using three different case studies.

### Benchmarking of mBARq using a previous *Salmonella* pathogenesis study extends the list of known fitness factors

To evaluate the performance of mBARq on a typical RB-TnSeq study, we re-analyzed a recent *Salmonella* Typhimurium mutagenesis screen [4]. In this screen, a small Tn5-based library of *Salmonella* mutants (∼2,000 strains, coverage of 392 insertions per Mb) was used to infect mice harboring a low complexity microbiome. The feces were collected on days 1, 2, 3, and 4 post infection (p.i.), and RB-TnSeq was used to identify *Salmonella* fitness factors impacting pathogenesis in this mouse model.

Taking the raw sequencing data from the experiment, we have re-created all the steps of the analysis using mBARq. First, a library map was generated. We identified 1,794 unique insertions, resulting in 858 genes with a transposon insertion within the coding region, and 773 genes with a disruption within the 5% to 95% percentile of the coding region. As expected, the insertions showed a largely random distribution along the chromosome (Fig. 2A). Next, we used mBARq to count the barcode abundances in the inoculum, as well as in the fecal samples at days 1 through 4 p.i. The count table was then uploaded to the mBARq web app for exploratory data analysis. We first generated a PCA plot (Fig. 2B), which showed a clear separation of samples based on days p.i., suggesting continuous selection against unfit mutants throughout the time course of infection. In addition, we observed high variability in samples collected on day 4 p.i., which was also reported in the original study. This is attributable to pronounced bottlenecks, which the host’s immune response inflicts upon the gut-luminal pathogen population [10].

**Figure 2:**
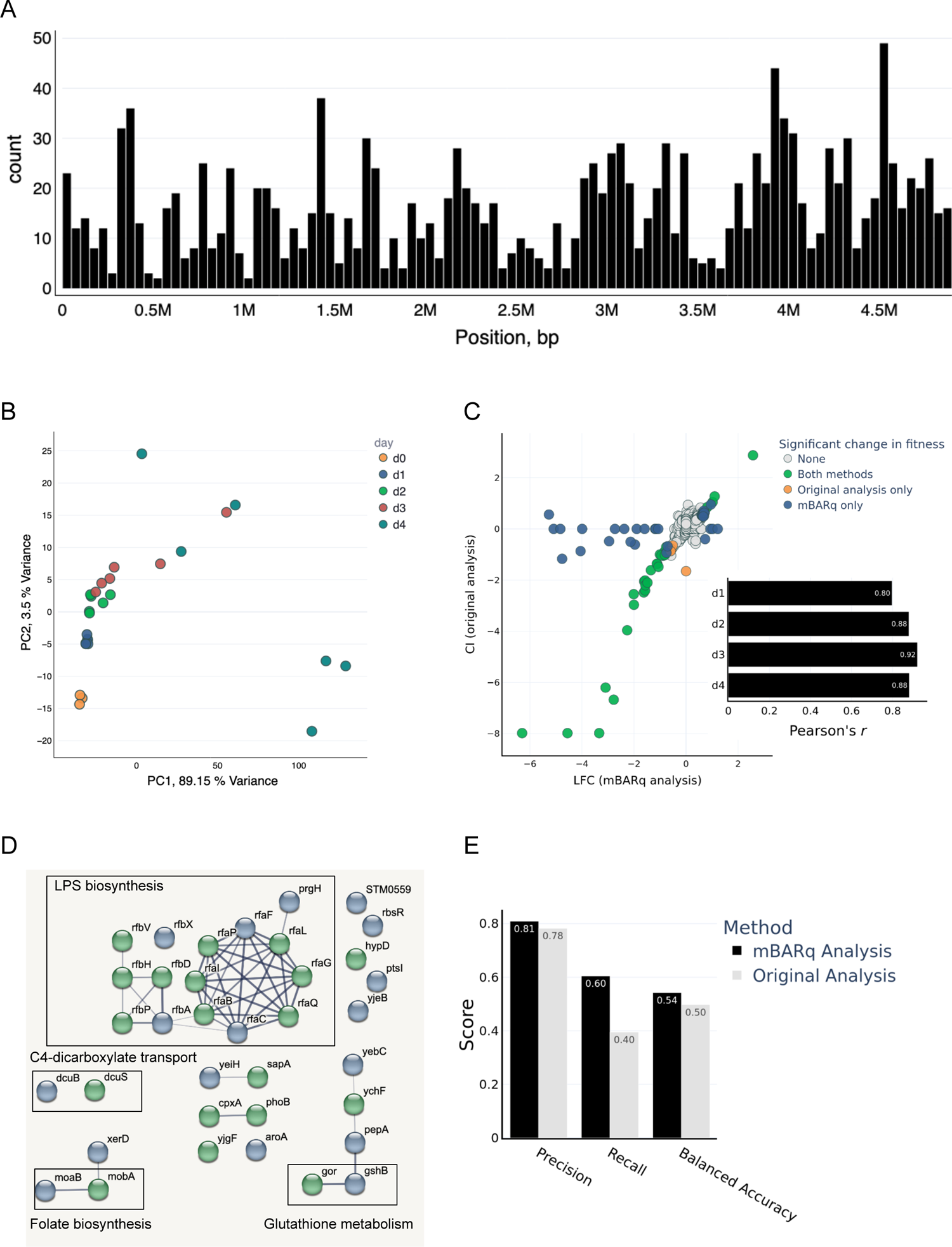
Benchmarking of mBARq using a previous *Salmonella* pathogenesis study extends the list of known fitness factors. **A.** Coverage histogram (bins size = 49 kbp) of *Salmonella* barcoded mutant library used by Nguyen *et al*. Generated with mBARq web app. **B.** PCA based on mutant strain abundances obtained from mouse fecal samples collected on different days p.i… The original inoculum is shown as d0. Generated with mBARq web app. **C.** Comparison of fitness effects reported in Nguyen *et al*. (show as log_2_ of competitive indices (CI)), and log_2_ fold changes (LFC) calculated by mBARq for day 1 p.i.. Genes were considered to have a significant change in fitness if absolute LFC (or log_2_(CI)) was greater than 0.6 and adjusted p-value was less than 0.05. Inset shows Pearson correlation between log_2_ CIs reported in the original analysis and LFC values calculated by mBARq for each day after infection. **D**. PPI network generated by STRING using genes with a fitness defect reported by mBARq on day 1 post infection. Original STRING-db network generated with mBARq web app was customized using a node coloring widget. Green: Significant change in fitness detected by both methods, blue: significant change detected by mBARq only. **E.** Benchmarking precision and recall of mBARq and previously published results using data from 28 clean gene KO strains competed 1:1 with WT *Salmonella* in the mouse [4].

To assess how well our analysis aligned with the previous one, we compared the fold changes computed by mBARq to the fitness estimates reported in the original publication. Overall, the results were highly concordant between the methods with the correlation between the data ranging between *r* = 0.8 and *r* = 0.92 (Fig. 2C). We further used mBARq to identify genes functioning as fitness factors on different days p.i.. To compare our results to the original study, we defined ‘hits’ (i.e., genes with a fitness defect) as genes with adjusted p-value < 0.05 and LFC (or log_2_ CI) < −0.6 for days 1 and 2 p.i.. mBARq was able to identify most of the hits previously identified in the study (26 out of 33). Moreover, mBARq reported 22 additional genes with significant defects in fitness on days 1 and/or 2 post infection. In order to validate these additional hits, we used the web app to submit hits identified by mBARq for day 1 p.i. to STRING and inspected the resulting PPI network (Fig 2D). The easy-to-use interface alleviated the need for dedicated support by a bioinformatician for performing this analysis and showed that many of the new hits belonged to the same biological process as the previously reported ones. For example, in addition to 10 genes belonging to the lipopolysaccharide biosynthesis pathway that were originally reported, mBARq identified 5 additional hits in the same pathway (Fig. 2D).

To further evaluate the precision of results produced by mBARq, we used experimental data from *in vivo* mutant competition experiments [4] for benchmarking. During the validation stage of the screen, Nguyen et al. (2020) generated site-directed knockout strains for 28 genes. These mutant strains were then competed one-on-one against a wild type strain in the same infection model, and fitness values for each gene were obtained on different days p.i. (total of 112 observations). We have used these experimental data to label genes as positive hits (defined as absolute LFC (or log_2_ CI) > 0.6 and p-value < 0.05), otherwise, they were considered negative. Using these data as the ‘ground truth’, we calculated precision, recall, and balanced accuracy (mean of recall values calculated separately for each class). The results show that mBARq has a comparable precision and accuracy (i.e., does not inflate false positives), and increased recall compared to the original method (Fig. 2E).

### mBARq extends functional insights in a previous *Shewanella amazonensis* metabolism study

To demonstrate the flexibility of mBARq, we applied it to the analysis of a large library (>380,000; 88,372 insertions per Mb) of *Shewanella amazonensis* mutants [5]. This library was constructed using a different *mariner*-based methodology and was used to investigate genes important for growth in the presence of different carbon sources. To this end, the *S. amazonensis* library was cultured in media containing various compounds as the only carbon source and sequencing data was generated as described above (Fig. 1). Subsequently, custom scripts and a statistical analysis model (hereafter referred to as Feba) was used to identify fitness factors for each growth condition.

Re-mapping the *S. amazonensis* library using mBARq identified a similar number of insertions as previously reported (380,770). Furthermore, the high coverage of transposon insertions across the chromosome was consistent with the published study (Fig. 3A). As above, we used mBARq to count barcode abundances and identify genes involved in carbon metabolism. Using an mBARq web app generated PCA plot, we observed a clear separation of the samples based on the carbon source and high consistency between the replicates (**Fig. S1**). Moreover, the counts and LFC calculated by mBARq were highly correlated with those reported in the original study (Fig. 3B). To further evaluate the concordance between mBARq and Feba, we compared the overlap between the hits as identified by the different methods. We defined hits for mBARq as described above (absolute LFC > 0.6 and adjusted p-value < 0.05), whereas Feba relies on a custom designed t-value, and defined hits as genes with |t| > 4. When looking at data across 25 different growth conditions, we found that the majority of the hits were identified by both methods. Across conditions, a mean of 49 hits (10%) were only identified by Feba and a mean of 188 hits (39%) only by mBARq (Fig. 3C).

**Figure 3:**
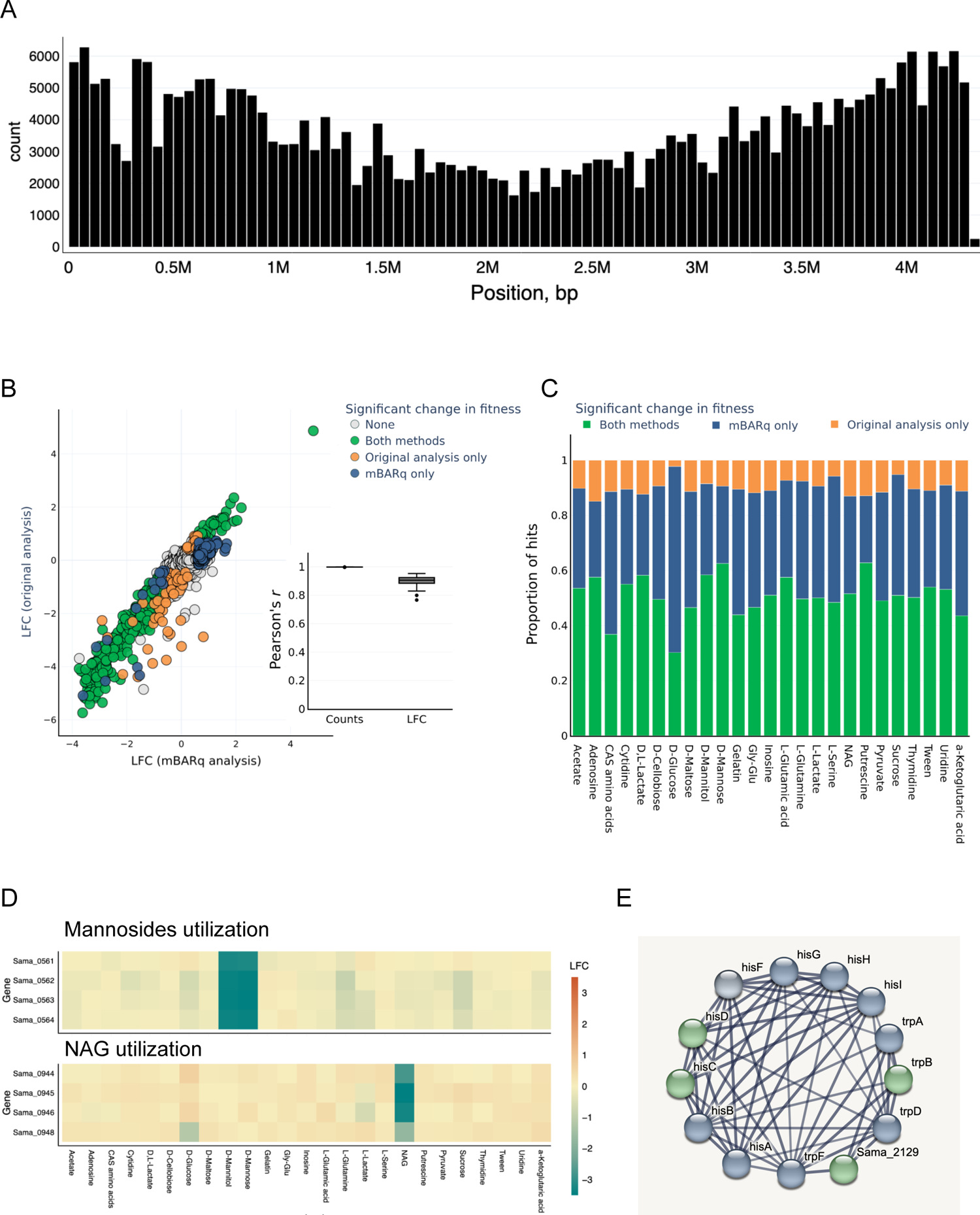
mBARq extends functional insights in a previous *Shewanella amazonensis* metabolism study. **A**. Coverage histogram of *S. amazonensis* barcoded mutant library used by Wetmore *et al*. Generated with mBARq web app. **B**. Comparison of LFCs reported in Wetmore *et al*., and LFCs calculated by mBARq for *S. amazonensis* cultured on Tween 20 as a sole carbon source. For mBARq, genes were considered to have a significant change in fitness if the absolute LFC was greater than 0.6 and the adjusted p-value was less than 0.05. Inset shows the Pearson correlation coefficient between counts and LFCs reported in the original analysis or calculated by mBARq for each culture condition (n=25). **C.** Proportions of fitness factors that were reported by both analysis methods, mBARq, or original analysis only for each culture condition. **D.** Heatmap of LFCs for genes predicted to play a role in mannoside utilization (top) or NAG utilization (bottom). Generated using mBARq web app. **E.** PPI network generated by STRING using genes with a fitness defect reported by mBARq in culture on D-Glucose. The PPI network was clustered using the MCL algorithm with default parameters. Only the largest cluster is shown. Original STRING-db network generated with mBARq web app was customized using a node coloring widget. Green: Significant change in fitness detected by both methods, blue: significant change detected by mBARq only.

To verify our results, we looked at the functional annotation of *Shewanella amazonensis* genes. Rodionov *et al.* [19] used comparative genomics to predict and experimentally validate genes responsible for carbon metabolism in different *Shewanella* species. This study delineated *S. amazonensis* genes that were important for N-acetyl glucosamine (NAG) and mannoside utilization. We hypothesized that mutations in these genes should result in a measurable fitness defect when cultured on NAG and D-Mannose, respectively. Using the mBARq web app, we generated LFC heatmaps for the genes of interest. We observed highly negative LFC for Sama_0944, Sama_0945, Sama_0946, and Sama_0948, when the library was cultured on NAG, but not other carbon sources. Furthermore, we observed a similar defect in Sama_0561, Sama_0562, Sama_0562, and Sama_0563, when *S. amazonensis* was cultured on either D-mannitol or D-mannose, confirming their role in mannoside utilization, and validating our analysis (Fig. 3D).

We also explored the PPI of the genes with a fitness defect in D-Glucose cultures, as mBARq reported a larger number of hits compared to Feba for this growth condition. We found that, while only *hisD* and *hisC* were identified in the original analysis, mBARq further identified *hisI*, *hisG*, *hisA*, *hisF*, *hisB* as having a fitness defect. Similarly, mBARq reported additional hits in the tryptophan biosynthesis pathway, which were missed by Feba (Fig. 3E). Finally, integration of mBARq analysis with STRING revealed the functional relevance of the gene Sama_2129, which lacks proper annotation in the *S. amazonensis* genome. A closer examination of Sama_2129 homologues revealed that it is likely encoding *trpE*, a key component of the tryptophan biosynthetic pathway. This finding highlights the power of our approach to improve the functional annotation of genes and thus help filling pathway gaps in the generation of genome-scale metabolic models. Furthermore, we hypothesized that the fitness defects of histidine metabolism mutants observed in D-Glucose culture should be alleviated during growth on Casamino acids. To compare histidine metabolism between these two conditions, we used the mBARq web app to integrate fitness data with the KEGG map of histidine metabolism and were able to confirm the differential requirement for histidine biosynthesis between these two culture conditions (**Fig S2**).

### mBARq recapitulates the evolutionary trajectories of barcoded isogenic *Escherichia coli* strains over 420 generations

Having demonstrated the precision and specificity of mBARq on RB-TnSeq experiments, we sought to demonstrate its utility beyond mutagenesis screens. Neutral genetic tags have become a powerful tool to decipher the dynamics of host-microbe interactions, as well as evolutionary trajectories of bacterial populations. While these studies were initially limited to a small number of barcoded tags, whose abundances could be quantified using qPCR, recent advances have simplified the generation of large barcode-tagged populations [20,21]. In one such study, the authors generated >400,000 distinct *E. coli* strains, each carrying a unique barcode inserted in a fitness-neutral location on the chromosome, using a method based on Tn7 transposon [20]. This barcoded *E. coli* population was repeatedly passaged in the presence of different antibiotic concentrations to better understand the evolutionary dynamics under antibiotic stress.

Here, we used the raw sequencing data from the original study to quantify the abundance of each barcoded strain in each of the bacterial passages. We then used the barcode count data to plot the abundance of individual strains across 420 bacterial generations (**Fig. 4A**). Our analysis results in evolutionary trajectories that are almost identical to those reported by the authors [20]. In addition, using data provided by the authors for the 20 most abundant lineages, we demonstrate the mean lineage frequencies, as well as the final frequencies (at generation 420), calculated by mBARq are virtually the same as those reported in the original study (**Fig. 4** **B, C**). Overall, this demonstrates mBARq’s broad applicability beyond barcoded mutagenesis experiments.

**Figure 4:**
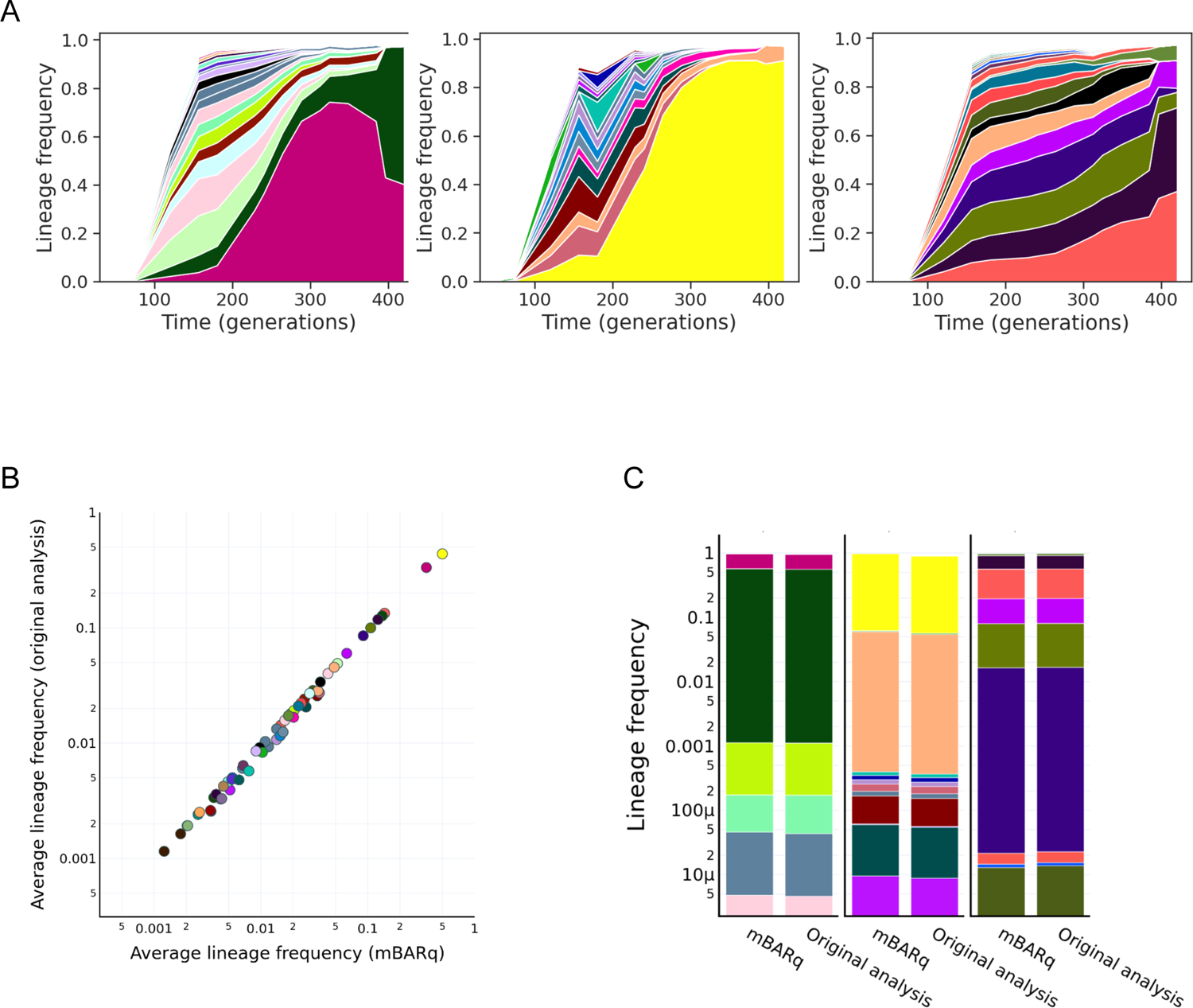
mBARq recapitulates the evolutionary trajectories of barcoded isogenic *E. coli* strains over 420 generations. **A.** Each panel shows the frequency trajectories for barcoded *E. coli* populations cultured without an antibiotic over 420 generations. Each colored band corresponds to a unique lineage, with its vertical width indicating its frequency at a particular time point. Only the 20 most abundant lineages are shown. **B**. Correlation between average frequencies for the 20 most abundant lineages from each of the replicates as calculated from mBARq generated counts and the original analysis. **C**. Final frequencies (i.e., at generation 420) of the 20 most abundant barcodes as calculated from mBARq counts and in the original analysis.

## DISCUSSION

Here we present mBARq, a versatile platform for the processing and analysis of barcoded sequencing data. We show that mBARq can readily accommodate different experimental setups and generate functional insights from RB-TnSeq and other barcode-based experiments, making these technologies accessible to a broad range of experimental scientists. With this in mind, we have developed documentation and walk-throughs along with test data that would allow non-expert users to recreate our analysis and to adapt mBARq for their own use [16].

By introducing a novel statistical methodology [12] to the analysis of barcoded sequencing data, mBARq not only reproduces previous results, but also identifies a larger number of significant hits in RB-TnSeq experiments. This increased sensitivity allows for the investigation of undetected fitness factors, with no significant loss in precision (i.e., higher rates of false positives). To the best of our knowledge, this is the first study that benchmarks RB-TnSeq analytical pipelines using knockout strains. In this regard, mBARq represents an improvement over existing methods, and highlights the importance of ‘ground truth’ data. The generation of additional ‘ground truth’ datasets, in particular ones containing more data on true negatives (i.e., genes with no fitness effect), will allow for better benchmarking and further development of statistical methods. In addition, the performance accuracy can also be improved by robust experimental design, such as increasing the number of replicates and including spike-in control barcodes to better estimate the variance in data.

mBARq was designed to be versatile and agnostic to the details of the library design or screening protocols. Thus, it is not only compatible with diverse transposon mutagenesis experimental setups, but also applicable to any barcoded sequencing data, including studies on site-specific mutants, as well as recently designed systems for genome editing in bacterial communities. One example is the DART (DNA-editing All-in-one RNA-guided CRISPR-Cas Transposase) system [22], which relies on barcoded transposon constructs, and should therefore be fully compatible with mBARq for analysis and visualization.

We showed that inspecting fitness data in the context of metabolic pathways or PPI networks often reveals phenotypes that are not apparent at the individual gene-level. In addition, we demonstrate that the re-analysis of published data using mBARq can also help to annotate genes that otherwise remain functionally uncharacterized. Finally, the integration of screening results with information stored in functional databases remains a common stumbling block for many researchers. Here, mBARq’s web app will empower biologists with little computational background to translate large amounts of screen data into biological insights.

## CONCLUSIONS

In conclusion, we show that mBARq contributes to the standardization and ease of data analysis for transposon insertion libraries, isogenic strain populations, and other barcoded sequencing data. The lack of user-friendly and flexible software for barcoded sequencing data analysis often results in researchers requiring bioinformatic expertise to analyze the data. In addition, variations in data processing and statistical methods often render cross-study comparisons difficult. In this study, re-analysis of public RB-TnSeq datasets with mBARq demonstrated the increased sensitivity of the tool and the user-friendly web app facilitated data interpretation and visualization to yield novel biological insights. Consequently, a tool such as mBARq is a major step forward towards the analysis, reuse, and cross-study comparisons of barcoded sequencing studies.

## METHODS

### Quality control of sequencing data

All sequencing data from the experiment were preprocessed using BBTools [23]. Specifically, adapters and potential contaminants (PhiX) were removed, and reads were quality-trimmed and filtered using BBDuk. The specific commands used are documented on our Methods in Microbiomics webpage [24] and in the mBARq documentation [16].

### mBARq Mapping

Mapping of the barcoded libraries was performed using *mbarq map* command. The specific commands used for each dataset can be found in mBARq documentation under Workflows [16]. During the mapping step, for each read, mBARq extracts the barcode and host sequence guided by user-provided specifications of barcode length and the end sequence of the transposon/construct used. If the identified host sequence is 20 nucleotides or longer, it is compared to the genome sequence using BLAST [25]. The BLAST results are filtered to keep only the matches with the highest bitscore. Analogous to the algorithm designed by Wetmore et al. [5], barcodes mapped to multiple locations (< 75% or reads map to primary location) are removed and the barcodes with an edit distance of < 2 and mapped to the same position are merged, with the more abundant barcoded assumed to be correct. Given an annotation file, mBARq also reports the closest feature of interest with attributes specified by the user, as well as how close the insertion is to the beginning/end of the feature (insertion percentile).

### mBARq Counting

Counting of the barcodes in each of the samples was performed using *mbarq count* command. The specific commands used for each dataset can be found in mBARq documentation under Workflows [16]. During this step, mBARq extracts barcode from each read guided by user-provided specifications of barcode length and the end sequence of the transposon/construct used. The frequency of each barcode is then summarized into a barcode count table. If specified by the user, the counts for barcodes below a specific edit distance can be merged, keeping the more abundant one, or the one that appears in the mapping file. The count tables from multiple samples can be merged using *mbarq merge*.

### mBARq Analysis

Analysis of the data was performed using *mbarq analyze* command. The specific commands used for each dataset can be found in mBARq documentation under Workflows [16]. The analysis step runs MAGeCK algorithm [12,13] to identify positively and negatively selected barcodes and genes, and consists of 4 steps (as described in the original publication): read count normalization, mean-variance modeling, barcode ranking, and gene ranking. For the experiments with control barcodes, *control* normalization method was chosen, otherwise the default *median* normalization was performed. Gene ranking was performed using MAGeCK’s robust rank aggregation (RRA) method.

### Using wildtype tagged strains as controls

When control barcoded strains were used in the experiments [4], mBARq also performed quality control on the samples, as described in Nguyen et al. Briefly, a linear regression is made with the dilution series of wild type control strains to verify that r^2^ > 0.8. If this is not the case, the sample is removed from the analysis. The minimum number of control barcodes required is 4.

---

**Figure S1:**
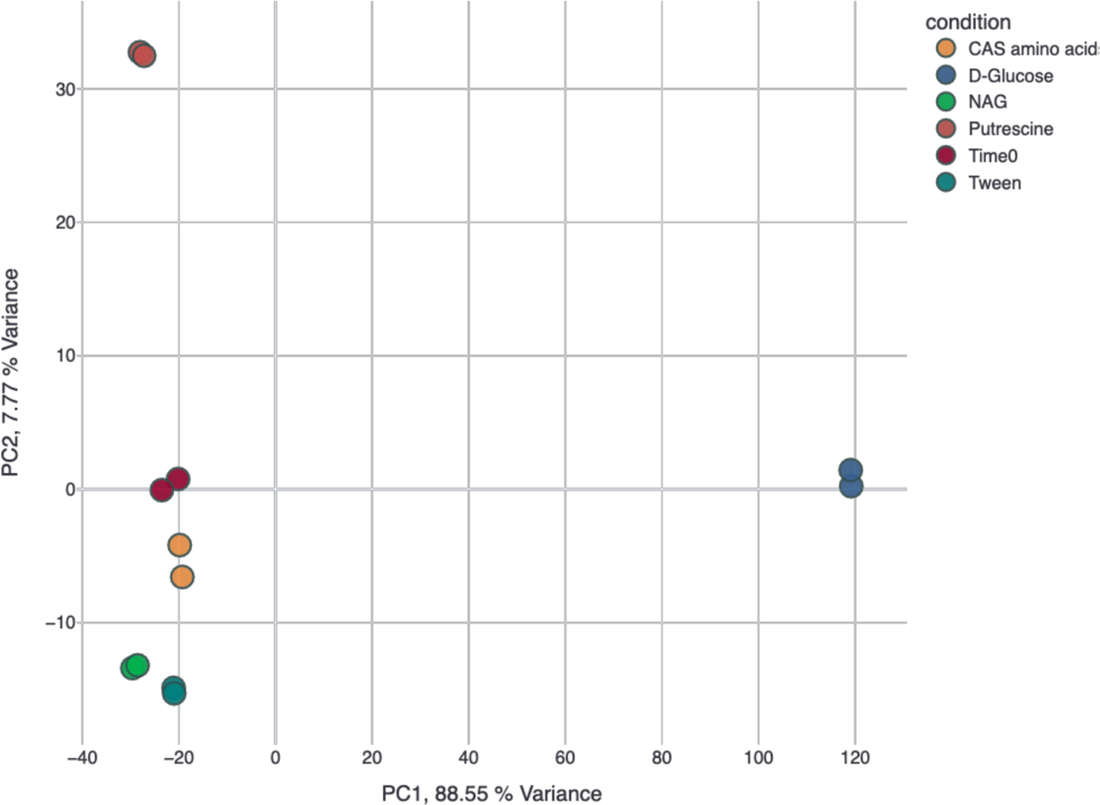
PCA based on mutant strain abundances obtained from *Shewanella amazonensis* cultured on different carbon sources. The original inoculum is shown as Time 0. Generated with mBARq web app.

**Figure S2:**
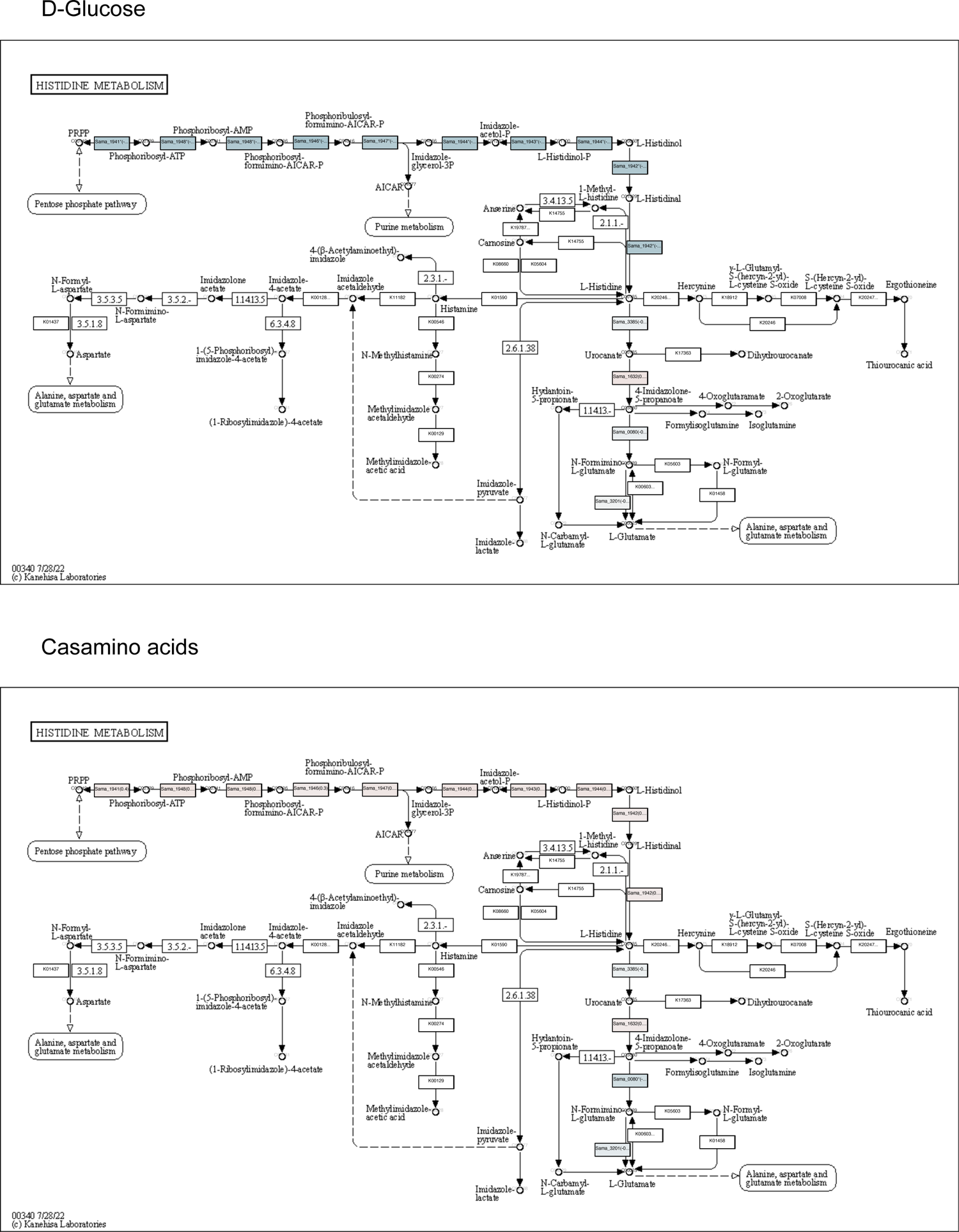
Fitness data for growth on D-Glucose and Casamino acids integrated with histidine metabolism pathway from KEGG database. Generated using mBARq web app. Blue indicates decreased fitness, and red indicates increased fitness. White boxes represent KOs not found in the genome.

## DECLARATIONS

### Ethics approval and consent to participate

Not applicable

### Consent for publication

Not applicable

### Availability of data and materials

All the data analysis for this manuscript was performed using Python and Jupyter notebooks, and the data and code to reproduce the figures are available at https://github.com/MicrobiologyETHZ/mbarq The mBARq web app is available at https://mbarq-app.herokuapp.com/ mBARq code and documentation are publicly available at https://github.com/MicrobiologyETHZ/mbarq Raw data used for the re-analysis is available as follows: Nguyen *et al*: ENA Project PRJEB63580; Wetmore *et al.*: https://genomics.lbl.gov/supplemental/rbarseq/; Jasinska *et al.*: SRA BioProject <u>PRJNA592371</u>

## Acknowledgements

ETH Zurich IT services and compute facilities.

## Author contributions

AS developed the tool and the web app, and designed and performed the analyses. LF contributed to the data analysis and app development. HJR and CF contributed to tool development. BN, BD, JV and WDH helped with technical clarifications and contributed to interpreting results. AS and SS wrote the manuscript with help from all the other authors. SS supervised the project. All authors read and approved the final manuscript.

